# RatesTools: a Nextflow pipeline for detecting *de novo* germline mutations in pedigree sequence data

**DOI:** 10.1101/2022.07.18.500472

**Authors:** Ellie E. Armstrong, Michael G. Campana

## Abstract

**Summary:** Here, we introduce *RatesTools*, an automated pipeline to infer *de novo* mutation rates from parent-offspring trio data of diploid organisms. By providing a reference genome and high-coverage, whole-genome resequencing data of a minimum of three individuals (sire, dam, offspring), RatesTools provides a list of candidate *de novo* mutations and calculates a putative mutation rate. RatesTools uses several quality filtering steps, such as discarding sites with low mappability and highly repetitive regions, as well as sites with low genotype and mapping qualities to find potential *de novo* mutations. In addition, RatesTools implements several optional filters based on *post hoc* assumptions of the heterozygosity and mutation rate of the organism. Filters are highly customizable to user specifications in order to maximize utility across a wide-range of applications.

**Availability:** RatesTools is freely available at https://github.com/campanam/RatesTools under a Creative Commons Zero (CC0) license. The pipeline is implemented in Nextflow (Di Tommaso et al. 2017), Ruby (http://www.ruby-lang.org), Bash (https://www.gnu.org/software/bash/), and R (R Core Team 2020) with reliance upon several other freely available tools. RatesTools is compatible with macOS and Linux operating systems.

**Supplementary Information:** We document RatesTools’ performance using published datasets in the online Supplementary information.

## 1. Introduction

Historically, germline mutation rates have been estimated primarily using neutral substitutions between species, often in combination with fossil information and predicted historical population sizes (Nachman and Crowell 2000). Alternatively, one can use mutation accumulation experiments, primarily suited for organisms traditionally grown in the lab with quick generation times and preferably with selfing capabilities (Fiston-Lavier et al. 2010; Zhu et al. 2014). Mutation rates have also been estimated using dominant disorders that rely on large counts of affected and unaffected individuals. Over time, improvements in the error rate of sequencing technology have made direct investigation of germline mutation rates using trios and pedigrees more feasible (Besenbacher et al. 2019; Campbell et al. 2021; Koch et al. 2019; Pfeifer 2017).

The calculation of an organism’s *de novo* germline mutation rate using a trio-based strategy is not without its issues. For one, when dealing with species of conservation concern, it can be difficult to obtain high-quality samples from verified trios, or across known pedigrees.

Secondarily, because *de novo* mutations are extremely rare, sequencing error, as well as error derived from mapping to repetitive or poorly assembled regions, can strongly contribute to error in mutation rate calculations (reviewed in Bergeron et al. 2022). Somatic mutations can also be difficult to parse from germline mutations, especially in certain tissue types (e.g., skin Martincorena et al. 2015), or based on the age (Cagan et al. 2022) or sex (Wilson Sayres and Makova 2011) of the individual.

Since the mutation rate has a profound impact on estimating divergence times, projecting historical population sizes, and building demographic models, more accurate estimations of mutation rate are needed. In order to estimate the *de novo* mutation rate directly from pedigree data, high-quality sites that can be confidently mapped against using short-read data must first be located, followed by filtering for genotype quality, which can differ based on the sequencer used as well as the depth of sequencing.

RatesTools provides the first customizable pipeline with visual and text outputs of various variant-call quality measures, allowing the user to adjust filters according to the data quality and depth in a convenient Nextflow (Di Tommaso et al. 2017) wrapper. It implements a number of different post-call filters, all of which can be customized to adjust for false positives. It also provides a detailed output of candidate single-nucleotide polymorphisms (SNPs) *de novo* mutations (DNMs), which can subsequently be verified empirically.

## 2. Implementation

RatesTools is built to take raw sequence data from parent-offspring trios and a reference fasta file as input and produce a set of candidate germline *de novo* mutations (DNMs). Currently, RatesTools is limited to diploid organisms, but the pipeline could be extended in the future to other ploidies. At various steps of the pipeline, summary statistics and graphs are produced to allow the user to adjust the pipeline’s filtering parameters based on the quality and depth of their input data.

See the Supplementary Text for a detailed description of the pipeline processes. An overview is presented in Figure 1. In brief, reads are mapped to the reference sequence using BWA-MEM (Li 2013). Alignments are finalized using SAMtools (Li et al. 2009), Picard (Broad Institute 2016), and the Genome Analysis Toolkit (GATK: McKenna et al. 2010). SNP calling is performed using GATK HaplotypeCaller. Users can also indicate which scaffolds are autosomal, if known. At this stage, graphs of GQ (genotype quality) and DP (depth) of all sites are produced for each individual, allowing the user to more easily select a cutoff for these filters.

**Figure 1:**
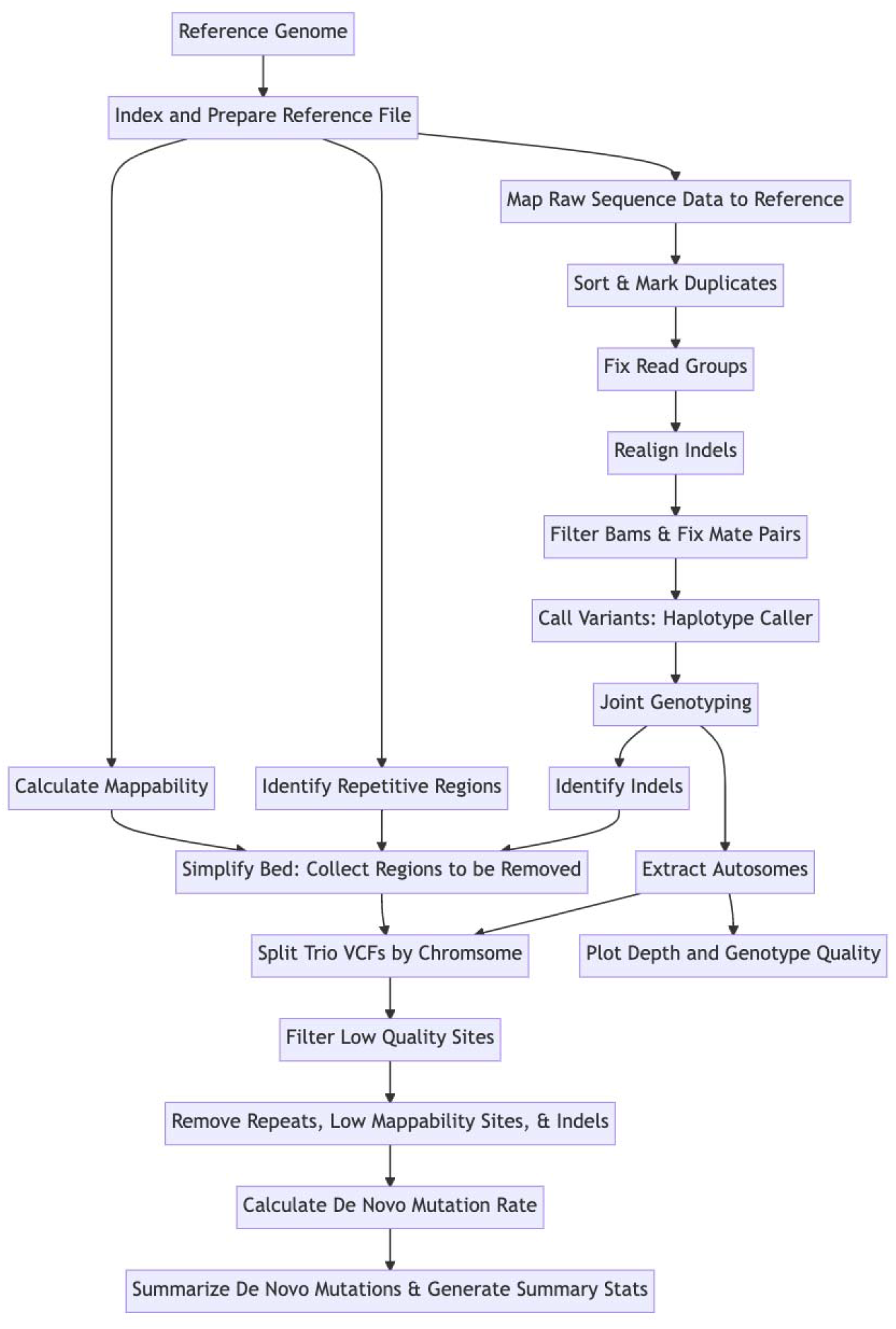
Schematic of RatesTools built using Mermaid v9.1.3 Live Editor.

After mapping, SNP calling, and optional autosomal-chromosome filtering, RatesTools performs site-level filtering using VCFtools (Danecek et al. 2011) and GATK. Finally, RatesTools removes repetitive elements, insertions and deletions, and sites identified as having low mappability, since these sites are likely to contain the most mapping or variant-calling related error. The sites remaining after the filtration steps are considered the ‘callable genome’.

After the determination of callable sites, candidate DNMs are identified using custom Ruby scripts. Additional filters based on previous studies (Besenbacher et al. 2019; Campbell et al. 2021; Koch et al. 2019; Pfeifer 2017; Venn et al. 2014) can be implemented by the user at this stage to reduce false positives. RatesTools then calculates a point estimate of the mutation rate. Optionally, RatesTools can perform block bootstrapping to estimate the 95% confidence interval of the mutation rate point estimate.

## 3. Application

To demonstrate the applicability of RatesTools, we used two pre-existing datasets for which the germline mutation rate has been calculated directly from sequence data, and candidate DNMs subsequently verified through PCR or genotyping. The first utilizes a wolf pedigree (*Canis lupus*: Koch et al. 2019) with one set of parents and three offspring (Supplementary Table S1). The second dataset consists of two chimpanzee pedigrees (*Pan troglodytes*: Venn et al. 2014; Supplementary Table S2). RatesTools identified candidate DNMs and calculated mutation rates that were comparable to the published literature (see *Supplementary Methods* for details). RatesTools identified all of the verified wolf DNMs, with a similar number of total candidates and fewer false positives. RatesTools yielded error-unadjusted point estimates that were slightly smaller (individual point estimates ranged between 3.7e-8 and 4.7e-8 mutations per base pair per generation) than those reported in Koch et al. (unadjusted rates between 5.1e-8 and 5.2e-8). RatesTools recovered 38 of 60 verified chimpanzee DNMs (63%), with the remaining sites either not being uniquely identified in the revised chimpanzee genome assembly or failing RatesTools quality filters (Table S9). Nevertheless, the chimpanzee mutation rate estimates across the six individuals tested (individual point estimates ranged between 1.6e-8 and 5.6e-8) were within an order of magnitude of the previously published value (1.2e-8).

## 4. Conclusions

RatesTools provides a framework for estimating *de novo* germline mutations from pedigree data using flexible filtering options that can be customized to sequence depth and quality. Implementation through a Nextflow wrapper provides convenient installation and deployment for the user.

## Acknowledgements

We thank D. Petrov and E. Hadly and members of both labs for helpful suggestions on early versions of the pipeline and J. Kelley for manuscript feedback. We also thank E. M. Koch and R. M. Schweizer for helpful input on the wolf dataset used to test the pipeline.

