## Supplementary File 1 for "RatesTools: a Nextflow pipeline for detecting *de novo* germline mutations in pedigree sequence data"

### Supplementary Text

#### Table of Contents

1. RatesTools Data Requirements
2. Installation and Dependencies
3. Pipeline Configuration and Execution
4. Tutorial and Test Dataset
5. Pipeline Processes

5a. Reference Genome Processing

5b. Read Mapping

5c. BAM Filtering

5d. Variant Calling

5e. De Novo Mutation Identification and Rate Calculation

6. Verification of Pipeline Using Pre-Existing Datasets

6a. Datasets

6b. Methods

6c. Results

6cI. Candidate Sites and Mutation Rates

6cII. Filtration Effects

6d. Discussion

6dI. Candidate Sites and Mutation Rates

6dII. Site Validation and Error Estimation

7. Future Directions
8. References
9. Supplementary Tables
10. Supplementary Figures

#### 1. RatesTools Data Requirements

RatesTools requires resequencing data for at least three individuals in a parent-offspring trio (sire, dam and offspring). For each individual in the trio, RatesTools needs a set of paired read files in FASTQ format. RatesTools assumes there is one forward and reverse read file per individual. If multiple read files exist per individual, these read files should be concatenated appropriately. Additionally, the end user must provide a reference genome file in FASTA format and, optionally, a list of chromosomes to retain in the final analysis.

#### 2. Installation and Dependencies

The RatesTools pipeline is implemented through the domain-specific language Nextflow v. 20.10.0 (Di Tommaso et al. 2017). It uses custom analysis scripts written in the Ruby v 2.6.3 (<http://www.ruby-lang.org>), Bash v.4.2.46(2)-release (<https://www.gnu.org/software/bash/>) and R v. 4.0.2 (R Core Team 2020) programming languages. RatesTools installation instructions are provided in the documentation (<https://github.com/campanam/RatesTools>). Installation of the RatesTools pipeline and custom scripts is automated using the provided Makefile.

RatesTools requires a number of freely available common bioinformatics packages. From the HTSlib package and libraries, RatesTools requires SAMtools, BCFtools, bgzip and tabix (Li et al. 2009; Danecek et al. 2021). RatesTools requires Picard (Broad Institute 2016), the Genome Analysis Toolkit (GATK: McKenna et al. 2010), and their dependency of Java 1.8 (<https://www.oracle.com/java/>). Other dependencies include BWA (Li 2013), Sambamba (Tarasov et al. 2015), VCFtools (Danecek et al. 2011), GenMap (Pockrandt et al. 2019), SeqAn (Reinert et al. 2017), BEDTools (Quinlan and Hall 2010), RepeatMasker (Smit et al. 1996) and RepeatModeler (Flynn et al. 2020). Within the R environment, RatesTools depends on the dplyr (Wickham et al., 2021), ggplot2 (Wickham 2016), tidyverse (Wickham et al. 2019) and data.table (Dowle and Srinivasan 2021) packages. Additionally, the RatesTools pipeline requires standard utilities (gzip, grep, awk, sed, zcat) provided by most Unix and Linux operating systems. To simplify installation and limit dependency conflicts, RatesTools can utilize modules specific to each process. Further facilitating deployment of the dependencies, Nextflow (Di Tommaso et al. 2017) has built-in support for Conda environments (Anaconda Inc., 2017) and containerization (e.g., Docker [<https://www.docker.com/>]).

#### 3. Pipeline Configuration and Execution

Analysis configuration is provided to RatesTools via a configuration file. The package includes a template configuration file (nextflow.config) that can be edited using a text processor. An interactive configuration script (configure.sh) is provided to assist analysis set-up. The script detects software installed on the local system and prompts the user to provide module files, paths to undetected files, and program options. We provide a detailed tutorial that explains run configuration (<https://github.com/campanam/RatesTools/blob/main/doc/tutorial.md>) and documentation for the custom RatesTools scripts (<https://github.com/campanam/RatesTools/blob/main/doc/pipeline_details.md>) to assist appropriate run set-up.

Given the wide-variety of computing architectures and operating systems, we cannot provide specific optimized configurations for all computing systems. However, the nextflow.config template has example profiles for a 'standard' local installation and one optimized for the SI/HPC Univa Grid Engine (UGE) computing cluster 'Hydra'. Nextflow (Di Tommaso et al. 2017) supports a wide variety of computing cluster architectures and cloud services, permitting these profiles to be adapted to most common systems.

After configuration, the pipeline is run by executing the primary RatesTools script (ratestools.nf) and specifying the configuration file and profile.

#### 4. Tutorial and Test Dataset

We provide a tutorial (<https://github.com/campanam/RatesTools/blob/main/doc/tutorial.md>) that explains RatesTools run configuration and executes the pipeline on a subset (chromosomes 1 and 2) of the wolf (*Canis lupus*: Koch et al. 2019) dataset described below. The subset data are available in the Smithsonian Figshare repository (DOI: 10.25573/data.20250288). The complete wolf datasets are available in the NCBI Sequence Read Archive (Accessions SRR1518530-SRR1518532, SRR9095635-SRR9095636,SRR9095638 from BioProjects PRJNA255370 (Fan et al. 2016) and PRJNA543877 (Koch et al. 2019). The reference chromosome sequences derive from the domestic dog (*Canis familiaris*) reference genome, build CanFam3.1 (Hoeppner et al. 2014). Users are encouraged to test the pipeline using this dataset to ensure that all commands are executing properly.

#### 5. Pipeline Analysis Stages and Processes

Here we provide a detailed summary of the pipeline analysis stages and processes (See Figure S1). We annotate where pipeline processes vary between GATK major builds (either GATK version 3 [GATK3] or version 4 [GATK4]). Further description of the individual Nextflow processes are provided in the online documentation (<https://github.com/campanam/RatesTools/blob/main/doc/pipeline_details.md>).

##### 5a. Reference Genome Processing

During this step, we prepare the reference genome sequence for downstream read mapping and genotyping and identify genomic regions prone to mismapping (repetitive regions and regions of low mappability). First, we index the reference genome and generate a sequence dictionary using BWA v. 0.7.17 (Li 2013) and SAMtools v. 1.9 (Li et al. 2009; Danecek et al. 2021). Next, repeat regions are identified first using RepeatMasker v. 4.0.9 (options ‘-gccalc -nolow’: Smit et al. 2013-2015) and a user-defined lineage-specific repeat library. Subsequently, RepeatModeler v. 2.0.1 (Flynn et al. 2020) is used to generate a species-specific repeat library for the reference sequence. This species-specific library is then used to annotate the repetitive regions in a second RepeatMasker run. The final file containing the coordinates of the repeat regions is then converted to BED file format using the custom script RM2bed.rb. Simultaneously, genomic regions with low mappability are identified with GenMap v. 1.2.0 (Pockrandt et al. 2019) using the ‘genmap index’ and ‘genmap map’ (options ‘-L 30 -E 2 -b’) commands. Sites with mappability < 1.0 are then filtered using a custom script (filterGM.rb).

##### 5b. Read Mapping

During this step, raw sequence data is aligned to the indexed reference sequence using BWA-MEM v. 0. 7.17 (Li 2013) and converted to sorted BAM format using SAMtools ‘view’ and ‘sort’ (Li et al. 2009). Duplicates are then marked using either Sambamba v. 0.7.1 ‘markdup’ (Tarasov et al. 2015) or Picard v. 2.23.8 ‘MarkDuplicates’ (Broad Institute 2016). Read groups are added and BAM indexes are built using the Picard ‘AddorReplaceReadGroups’ and ‘BuildBamIndex’ commands respectively. If GATK3 is used, indels are then realigned using the ‘ReAlignerTargetCreator’ and ‘IndelRealigner’ (option ‘--filter_bases_not_stored’) commands. GATK4 analyses use the ‘LeftAlignIndels’ command to left-align indels.

##### 5c. BAM Filtering

In this step, we filter BAMs using the GATK ‘PrintReads’ command with the objective of minimizing reads that either have poor sequence quality or mapping scores. For GATK3 we use the ‘BadCigar’, ‘DuplicateRead’, ‘FailsVendorQualityCheck’, ‘HCMappingQuality’, ‘MappingQualityUnavailable’, ‘NotPrimaryAlignment’, and ‘UnmappedRead’ read filters along with the options ‘--filter_bases_not_stored’ and ‘--filter_mismatching_base_and_quals’. For GATK4, we use the ‘GoodCigarReadFilter’, ‘NotDuplicateReadFilter’, ‘PassesVendorQualityCheckReadFilter’ , ‘MappingQualityReadFilter’, ‘MappingQualityAvailableReadFilter’, ‘PrimaryLineReadFilter’, ‘MappedReadFilter’, ‘NotOpticalDuplicateReadFilter’, and ‘ProperlyPairedReadFilter’ read filters. After filtering, we correct mate pair tags using Picard ‘FixMateInformation’ (option ‘ADD_MATE_CIGAR=true’) and build a BAM index using Picard ‘BuildBamIndex’.

##### 5d. Variant Calling

At this step we generate individual variant calls for all positions of the genome. This step is crucial, since the number of callable sites will strongly factor into our eventual mutation rate calculation. First, we call all sites for each individual using GATK ‘HaplotypeCaller’. For GATK3, we apply the options ‘-A DepthPerSampleHC -A Coverage -A HaplotypeScore -A StrandAlleleCountsBySample -ERC GVCF -out_mode EMIT_ALL_SITES’, while for GATK4, we use ‘-ERC GVCF -G StandardAnnotation -G AS_StandardAnnotation’. Subsequent to HaplotypeCaller, joint-genotyping is performed to generate final all-sites VCF files for all members of the cohort. For GATK3, this process uses ‘GenotypeGVCFs’ (option ‘--includeNonVariantSites’). For GATK4, this process first uses ‘CombineGVCFs’ (option ‘--convert-to-base-pair-resolution’) and then ‘GenotypeGVCFs’ (option ‘--include-non-variant-sites’).

Following joint-genotyping, we generate a BED file annotating the location of all indels and excluding a set number of base pairs (bp) up- and downstream of each indel using a custom script (indels2bed.rb). After indel discovery, we (optionally) retain only sites from chromosomes provided in the chromosome list. Next, we extract depth (DP) and genotype quality (GQ) statistics from all retained variable sites and plot their distributions using the R packages dplyr v.1.0.7 (Wickham et al. 2021), ggplot2 v.3.3.5 (Wickham 2016), tidyverse v. 1.3.1 (Wickham et al. 2019) and data.table v. 1.14.2 (Dowle and Srinivasan 2021). We then divide the chromosomal-filtered all-sites VCF into individual trio (one offspring and its two parents) and split the resulting files by chromosome for parallelization.

##### 5e. Variant Filtering

Each split VCF is then filtered at the site-level using VCFtools v. 0.1.16 (Danecek et al. 2011) and GATK (McKenna et al. 2010). Filter settings are entirely user-customizable, although we recommend that end users restrict their analyses to monoallelic and biallelic sites. Following site-level filters, low-mappability regions, repetitive regions, and regions around indels are removed using BEDTools v. 2.28.0 (Quinlan and Hall 2010), BCFtools (Li et al. 2009; Danecek et al. 2021), VCFtools, and custom scripts. After each filtration step, the number of retained sites is logged using VCFtools, BCFtools and custom scripts.

##### 5f. De Novo Mutation Identification and Mutation Rate Calculation

Following site- and region-filtering, DNM candidates are identified from each VCF using a custom script (calc_denovo_mutation_rate.rb). We assume that the sequencing error has been removed via the previous site-level filters. Since false positives are pervasive in current trio analyses (e.g. Koch et al. 2019), we have provided a number of filtration options that can be applied at this time to reduce noise. These include an option to treat the presence of any copies of an allele as a true allele (even if not called by GATK: option ‘--minAD1’) and an option to discard alleles below a specified frequency (option ‘--minAF’). When these two options are both specified, this produces maximally conservative behavior as the ‘--minAD1’ option is applied to the parents (where the most likely source of deviation from the expected 50%:50% allelic ratio is allelic dropout) and the ‘--minAF’ filter to the offspring (where the most likely source of allelic ratio deviation is somatic mutation). Restricting to sites where both parents are homozygous (e.g. Besenbacher et al. 2019; Campbell et al. 2021; Koch et al. 2019; Pfeifer 2017; Venn et al. 2014) can be performed using the ‘--parhom’ option. Filtering using the Koch et al. (2019) DNp statistic, derived from Ramu et al. (2013), can be performed using the option ‘--kochDNp’. Alternatively, the DNp statistic can be applied *post-hoc* to the RatesTools output using the kochDNp.rb script. Following candidate DNM identification, calc_denovo_mutation_rate.rb calculates a point estimate of the mutation rate using the formula (where μ is the mutation rate, *N* is the number of candidate DNMs, and *CG* is the callable genome in bp: Bergeron et al. 2022):

μ=*N*/(2×*CG)*

RatesTools does not apply a correction for false negatives or false positives (Bergeron et al. 2022; Koch et al. 2019; Besenbacher et al. 2019; Pfeifer 2017; Campbell et al. 2021) as estimating error rates typically requires additional empirical (e.g. Koch et al. 2019; Venn et al. 2014) or simulation analyses (e.g. Campbell et al. 2021). Optionally, block bootstrapping can be performed to estimate the confidence interval for the mutation rate estimates. RatesTools provides two estimates: one including all observed DNMs (single-forward mutations, double-forward mutations [forward mutations on both copies of sites], and back mutations) and one including only single-forward mutations (since recurrent DNMs at a single site are very unlikely: Venn et al. 2014). Finally, the individual chromosome-level results are combined using the summarize_denovo.rb script. The number of retained sites after each filtration step is tabulated and the number of each single-forward DNM substitution type (e.g. C → T, G → C) is calculated using the dnm_summary_stats.rb script.

#### 6. Verification of Pipeline Using Pre-Existing Datasets

##### 6a. Datasets

In order to validate the pipeline, we tested it on two pre-existing datasets of wolf (*Canis lupus*; Koch et al. 2019) and chimpanzee (*Pan troglodytes*: Venn et al. 2014) pedigrees. For the wolves, we analyzed a subset of the individuals in the paper, while for the chimpanzees, we analyzed all individuals, but as two separate pedigrees (details can be found in Supplementary Tables S1 & S2). In order to be consistent with the coordinate system used in Koch et al. (2019), we used Canfam3.1 (NCBI Accession: GCF_000002285.3: Hoeppner et al. 2014) as the wolf reference genome. For the chimpanzee dataset, we used assembly Pan_tro 3.0 (NCBI Accession: GCA_000001515.5), which is an updated version of the panTro3 (NCBI Accession: GCA_000001515.3: Kuderna et al. 2017) assembly used in Venn et al. (2014). To identify the verified DNM candidates from Venn et al. in the revised assembly, we mapped the 51 bp sequence context of each SNP to the Pan_tro 3.0 sequence using BLAST v. 2.10.1 (Camacho et al. 2009) and retained the highest-quality, complete-length match for each sequence. Specifically, we used *blastn* with flags ‘-perc_identify 95’ and ‘-evalue 1e-10’.

##### 6b. Methods

Analyses were performed using RatesTools under the default settings described above using GATK3 for comparability with the previous publications. Repetitive genomic regions were annotated using the ‘Canis lupus’ and ‘Pan troglodytes’ RepeatMasker libraries (Smit et al. 2013-2015). We restricted our analyses to scaffolds that were assigned to autosomes for each species. Subsequent to the removal of non-autosomal contigs and scaffolds, we filtered sites for depth (15 ≤ DP ≤ 125; see Table S3 and Table S4), genotype quality (GQ ≥ 40; see Table S3 and Table S4), an allele count of 1 or 2 (--min-alleles 1 and --max-alleles 2), and excluded sites with missing data. We then filtered sites with quality by depth score (QD) < 4.0, Fisher strand bias (FS) > 60.0, root mean square mapping quality across all samples (MQ) < 40.0 , symmetric-odds-ratio-tested strand bias (SOR) > 3.0, read position rank sum test (ReadPosRankSum) < 15 or mapping quality rank sum test (MQRankSum < -2). We then removed five bp up- and downstream of all identified indels in addition to low-mappability and repetitive regions. Given the sequencing depth (18–46×; Table S3 and Table S4) of the datasets, we identified candidate DNMs using conservative parameters, requiring parental homozygosity (option ‘--parhom’), discarding sites where parents had evidence of uncalled alleles (option ‘--minAD1’), and requiring that candidate offspring DNM alleles had a minimum frequency of 0.3 (option ‘--minAF 0.3). We applied the Koch et al. (2019) DNp statistic using the filter settings described there (mutation rate of 1e-6, heterozygosity of 0.008, and a DNp cut-off of 0.3). We block-bootstrapped mutation rate estimates using 100 bootstrap replicates, a window size of 100,000 bp, a window-step size of 50,000 bp, and a minimum of 10 windows to retain a chromosome. As in the previous studies, we only considered single-forward mutations as candidate DNMs.

We then compared the DNM candidates from our pipeline to the Sanger-verified candidates from Koch et al. (2019) and the genotyped variants from Venn et al. (2014). We tabulated the number of differences between our candidate mutations and the candidate sites from each study. Finally, to explore the effects of the various filters, we tabulated the numbers of candidate DNM sites retained after each filter was applied to the wolf dataset.

##### 6c. Results

###### 6cI. Candidate Sites and Mutation Rates

The wolf dataset yielded between 1.23 and 1.27 Gb of callable sites out of the approximately 2.39 Gb genome after filtering (Table S5). This is approximately 8% more of the genome than previously examined by Koch et al. (2019) (1.04Gb). For the wolf pedigree, RatesTools identified 94, 104, and 119 candidate DNM’s across the three offspring, respectively (Table S5). Comparatively, Koch et al. identified 109, 106, and 108 candidate DNMs for these individuals. Of those verified via Sanger sequencing, RatesTools identified all true positives and a total of 3/29 false positives identified by the original analysis (Table S8). This yielded a final error-unadjusted point mutation rate calculation of the wolves of between 3.7e-8 and 4.7e-8 mutations per base pair per generation for each individual. For comparison, Koch et al.’s unadjusted mutation rate is between 5.1e-8 and 5.2e-8 mutations per base pair per generation.

The chimpanzee data yielded between 1.43 and 1.65 Gb of callable sites out of the approximately 3.2 Gb genome after filtering (Table S6 and Table S7). This is substantially fewer than those evaluated by Venn et al. (2014), who evaluated 2.47 Gb of the approximately 3.2Gb genome. Pedigree 1 (Table S6) yielded slightly more callable sites across individuals than pedigree 2 (Table S7), likely because pedigree 1 was sequenced at higher coverage than pedigree 2 (Table S4).

For the three chimpanzee individuals in pedigree 1, RatesTools identified a total of 195, 154, and 192 candidate DNMs respectively (Table S6), while for the three individuals from pedigree 2, RatesTools identified 47, 79, and 152 candidate DNMs respectively (Table S7). For each of the chimpanzees that were validated by genotyping in Venn et al. (2014), we identified 24/35 and 14/25 true positive DNMs respectively (Table S9). Upon further investigation, we found that many of the candidates not identified by RatesTools were removed by various filters (see Table S9). Six of these non-identified candidates were found in panTro3 contexts that have multiple matches in the revised Pan_tro 3.0 genome, while another four were not recovered as SNP sites. Across the chimpanzee individuals, we calculated an error-unadjusted mutation rate of between 1.6e-8 and 5.6e-8 mutations per base pair per generation.

###### 6cII. Filtration Effects

Table S10 lists the number of DNM candidates observed after each filter. The most critical filters were the site-quality and the low-quality region filters which reduced the number of candidate single-forward DNMs by 22–30-fold each. The allele count filters (options ‘--minAD1’ and ‘--minAF 0.3’) also had a strong effect, reducing the number of candidates by 4.5–11-fold. While the parental homozygosity filter (option ‘--parhom’) only directly reduced the number unlikely recurrent DNMs, it also contributed to the reduction of candidates from the allele count filters as the homozygosity filter was responsible for removal of parental heterozygotes call homozygous by GATK. Overall, filtering reduced the number of candidates from a mean of 667,195 to a mean of 106, a reduction of 99.98%.

##### 6d. Discussion

###### 6dI. Candidate Sites and Mutation Rates

We validated the RatesTools pipeline using two existing whole-genome sequence datasets, consisting of pedigrees of wolves (Koch et al. 2019) and chimpanzees (Venn et al. 2014). Our results demonstrated that RatesTools performs well across various taxa and of datasets with varying depth and quality. Using tailored filtering based on the depth and quality of each of the datasets, we obtained error-unadjusted point estimates of the mutation rates of 3.7e-8 and 4.7e-8 mutations per base pair per generation for wolves and 1.6e-8 and 5.6e-8 for chimpanzees.

While Koch et al. (2019) report an error-adjusted mutation rate of 4.5e-9 for the wolf dataset, their unadjusted rates (5.1e-8 to 5.2e-8) are somewhat higher (11–-38%) than our estimates, mostly due to having recovered less of the callable genome. RatesTools also retained fewer false positives than the original wolf study (Table S8), while identifying similar numbers of total candidates and all of the empirically confirmed DNM sites. However, it is possible that we identified different, but similar numbers of false positives, making our error-adjusted mutation rate comparable with the previous study.

The mutation rate calculated by RatesTools for the chimpanzees is within an order of magnitude of the previous mutation rate estimate from Venn et al. (2014), despite RatesTools identifying more candidate sites. These differences could be attributable to a number of different steps taken in Venn et al. (2014) and not in RatesTools, and vice versa. Most importantly, their pipeline includes a manual curation step, but none of the sites removed by manual curation were queried for verification by genotyping to confirm that they were not actually DNMs. Additionally, their identified false positive site is not marked, so it was not possible for us to incorporate this into our results. The study also employs a different variant calling tool and filtering thresholds, all of which can lead to differences in candidate sites. A comprehensive comparison of various filtering strategies for DNM discovery is discussed in Bergeron et al. (2022). Notably, identified DNM candidates were dependent on the reference genome assembly version, despite using the same original sequencing data (Table S9).

###### 6dII. Site Validation and Error Estimation

Variant filtering is critical to reduce false positive DNM candidates, but introduces biases and assumptions about the underlying data. It is clear that empirical verification through direct genotyping or sequencing methods is a crucial step to both reduce and estimate the number of false positives. RatesTools provides a viable pipeline for generating a relatively small number of candidate sites that could be queried to estimate a mutation rate. Currently, verification of candidate DNMs is typically performed using polymerase chain reaction and Sanger sequencing (e.g., Koch et al. 2019). However, designing and testing primer sets for hundreds to thousands of candidate loci is an onerous and expensive task. We propose that a large number of candidate DNMs would be more effectively verified using hybridization capture (Gnirke et al. 2009) as thousands of candidates could be assayed simultaneously and cost-effectively sequenced on a current-generation sequencer. Moreover, the ability to genotype large numbers of candidates simultaneously would reduce the need for stringent candidate filtration steps that introduce strong experimenter biases. RatesTools outputs a final VCF of candidate DNMs that can be input directly into BaitsTools (Campana 2018) for bait generation.

#### 7. Future Directions

The RatesTools pipeline is highly flexible and readily expandible. Potential future revisions include support for non-diploid organisms and chromosomes (e.g., haplodiploidy) and automated estimation of the false negative rate (e.g. using simulated mutations: Campbell et al. 2021). The Nextflow processes used in the pipeline are highly modular, permitting the addition or replacement of component tools without the need to rewrite the entire codebase. For instance, alternative genotypers such as BCFtools (Danecek et al. 2021) or ANGSD (Korneliussen et al. 2014) could be substituted for GATK.

Camacho, Christiam, George Coulouris, Vahram Avagyan, Ning Ma, Jason Papadopoulos, Kevin Bealer, and Thomas L. Madden. 2009. “BLAST+: Architecture and Applications.” *BMC Bioinformatics* 10 (December): 421.

Campana, Michael G. 2018. “BaitsTools: Software for Hybridization Capture Bait Design.” *Molecular Ecology Resources* 18 (2): 356–61.

Campbell, C. Ryan, George P. Tiley, Jelmer W. Poelstra, Kelsie E. Hunnicutt, Peter A. Larsen, Hui-Jie Lee, Jeffrey L. Thorne, Mario Dos Reis, and Anne D. Yoder. 2021. “Pedigree-Based and Phylogenetic Methods Support Surprising Patterns of Mutation Rate and Spectrum in the Gray Mouse Lemur.” *Heredity* 127 (2): 233–44.

Danecek, P., A. Auton, G. Abecasis, C. A. Albers, E. Banks, M. A. DePristo, R. E. Handsaker, et al. 2011. “The Variant Call Format and VCFtools.” *Bioinformatics*. https://doi.org/[10.1093/bioinformatics/btr330](http://dx.doi.org/10.1093/bioinformatics/btr330).

Danecek, Petr, James K. Bonfield, Jennifer Liddle, John Marshall, Valeriu Ohan, Martin O. Pollard, Andrew Whitwham, et al. 2021. “Twelve Years of SAMtools and BCFtools.” *GigaScience* 10 (2). https://doi.org/[10.1093/gigascience/giab008](http://dx.doi.org/10.1093/gigascience/giab008).

Di Tommaso, Paolo, Maria Chatzou, Evan W. Floden, Pablo Prieto Barja, Emilio Palumbo, and Cedric Notredame. 2017. “Nextflow Enables Reproducible Computational Workflows.” *Nature Biotechnology* 35 (4): 316–19.

Dowle, Srinivasan, and Gorecki. n.d. “Data. Table: Extension of ‘Data. Frame’. R Package Version 1.12. 8.” *Manual de Referencia En Mejores Prácticas de Gestión de Datos Oceánicos*.

Fan, Zhenxin, Pedro Silva, Ilan Gronau, Shuoguo Wang, Aitor Serres Armero, Rena M. Schweizer, Oscar Ramirez, et al. 2016. “Worldwide Patterns of Genomic Variation and Admixture in Gray Wolves.” *Genome Research*. https://doi.org/[10.1101/gr.197517.115](http://dx.doi.org/10.1101/gr.197517.115).

Fiston-Lavier, Anna-Sophie, Nadia D. Singh, Mikhail Lipatov, and Dmitri A. Petrov. 2010. “Drosophila Melanogaster Recombination Rate Calculator.” *Gene*. https://doi.org/[10.1016/j.gene.2010.04.015](http://dx.doi.org/10.1016/j.gene.2010.04.015).

Flynn, Jullien M., Robert Hubley, Clément Goubert, Jeb Rosen, Andrew G. Clark, Cédric Feschotte, and Arian F. Smit. 2020. “RepeatModeler2 for Automated Genomic Discovery of Transposable Element Families.” *Proceedings of the National Academy of Sciences of the United States of America* 117 (17): 9451–57.

Hoeppner, Marc P., Andrew Lundquist, Mono Pirun, Jennifer R. S. Meadows, Neda Zamani, Jeremy Johnson, Görel Sundström, et al. 2014. “An Improved Canine Genome and a Comprehensive Catalogue of Coding Genes and Non-Coding Transcripts.” *PloS One* 9 (3): e91172.

Broad Institute. 2016. “Picard Tools.”

Koch, Evan, Rena M. Schweizer, Teia M. Schweizer, Daniel R. Stahler, Douglas W. Smith, Robert K. Wayne, and John Novembre. 2019. “De Novo Mutation Rate Estimation in Wolves of Known Pedigree.” *Molecular Biology and Evolution*, July. https://doi.org/[10.1093/molbev/msz159](http://dx.doi.org/10.1093/molbev/msz159).

Kuderna, Lukas F. K., Chad Tomlinson, Ladeana W. Hillier, Annabel Tran, Ian T. Fiddes, Joel Armstrong, Hafid Laayouni, et al. 2017. “A 3-Way Hybrid Approach to Generate a New High-Quality Chimpanzee Reference Genome (Pan_tro_3.0).” *GigaScience* 6 (11): 1–6.

[Li, Heng. 2013. “Aligning Sequence Reads, Clone Sequences and Assembly Contigs with BWA-MEM.” *arXiv [q-bio.GN]*. arXiv.](http://paperpile.com/b/oqYMtK/4EWN) <http://arxiv.org/abs/1303.3997>.

Li, Heng, Bob Handsaker, Alec Wysoker, Tim Fennell, Jue Ruan, Nils Homer, Gabor Marth, Goncalo Abecasis, Richard Durbin, and 1000 Genome Project Data Processing Subgroup. 2009. “The Sequence Alignment/Map Format and SAMtools.” *Bioinformatics*  25 (16): 2078–79.

Martincorena, Iñigo, Amit Roshan, Moritz Gerstung, Peter Ellis, Peter Van Loo, Stuart McLaren, David C. Wedge, et al. 2015. “Tumor Evolution. High Burden and Pervasive Positive Selection of Somatic Mutations in Normal Human Skin.” *Science* 348 (6237): 880–86.

McKenna, Aaron, Matthew Hanna, Eric Banks, Andrey Sivachenko, Kristian Cibulskis, Andrew Kernytsky, Kiran Garimella, et al. 2010. “The Genome Analysis Toolkit: A MapReduce Framework for Analyzing next-Generation DNA Sequencing Data.” *Genome Research* 20 (9): 1297–1303.

Nachman, M. W., and S. L. Crowell. 2000. “Estimate of the Mutation Rate per Nucleotide in Humans.” *Genetics* 156 (1): 297–304.

Pfeifer, Susanne P. 2017. “Direct Estimate of the Spontaneous Germ Line Mutation Rate in African Green Monkeys.” *Evolution*. https://doi.org/[10.1111/evo.13383](http://dx.doi.org/10.1111/evo.13383).

Pockrandt, Christopher, Mai Alzamel, Costas S. Iliopoulos, and Knut Reinert. 2019. “GenMap: Fast and Exact Computation of Genome Mappability.” *Cold Spring Harbor Laboratory*. https://doi.org/[10.1101/611160](http://dx.doi.org/10.1101/611160).

Quinlan, Aaron R., and Ira M. Hall. 2010. “BEDTools: A Flexible Suite of Utilities for Comparing Genomic Features.” *Bioinformatics*  26 (6): 841–42.

Ramu, A., Noordam, M. J., Schwartz, R. S., Wuster, A., Hurles, M. E., Cartwright, R. A., & Conrad, D. F. (2013). DeNovoGear: de novo indel and point mutation discovery and phasing. *Nature methods*, *10*(10), 985-987.

Reinert, Knut, Temesgen Hailemariam Dadi, Marcel Ehrhardt, Hannes Hauswedell, Svenja Mehringer, René Rahn, Jongkyu Kim, et al. 2017. “The SeqAn C++ Template Library for Efficient Sequence Analysis: A Resource for Programmers.” *Journal of Biotechnology* 261 (November): 157–68.

Ryan Campbell, C., George P. Tiley, Jelmer W. Poelstra, Kelsie E. Hunnicutt, Peter A. Larsen, Mario dos Reis, and Anne D. Yoder. 2019. “Pedigree-Based Measurement of the de Novo Mutation Rate in the Gray Mouse Lemur Reveals a High Mutation Rate, Few Mutations in CpG Sites, and a Weak Sex Bias.” *bioRxiv*. https://doi.org/[10.1101/724880](http://dx.doi.org/10.1101/724880).

Smit, Arian F. A., Robert Hubley, and P. Green. 1996. “RepeatMasker.”

Tarasov, Artem, Albert J. Vilella, Edwin Cuppen, Isaac J. Nijman, and Pjotr Prins. 2015. “Sambamba: Fast Processing of NGS Alignment Formats.” *Bioinformatics*  31 (12): 2032–34.

R Core Team and Others. 2020. “R: A Language and Environment for Statistical Computing. R Foundation for Statistical Computing.”

Venn, O., I. Turner, I. Mathieson, and N. de Groot. 2014. “Strong Male Bias Drives Germline Mutation in Chimpanzees.” <https://science.sciencemag.org/content/344/6189/1272.abstract?casa_token=i06j4rUcFAIAAAAA:G-Tc7-1ehTfCwqJsZxOVZoxq-einDtS21zi_sAUlQH2537FL2lwmv0VeurIjsEssQuLn6myz9fIqCw>.

Wickham, François, Henry, and Müller. n.d. “Dplyr: A Grammar of Data Manipulation.” *R Package Version 0.4*.

Wickham, Hadley. 2011. “Ggplot2.” *Wiley Interdisciplinary Reviews. Computational Statistics* 3 (2): 180–85.

Wickham, Hadley, Mara Averick, Jennifer Bryan, Winston Chang, Lucy McGowan, Romain François, Garrett Grolemund, et al. 2019. “Welcome to the Tidyverse.” *Journal of Open Source Software* 4 (43): 1686.

Wilson Sayres, Melissa A., and Kateryna D. Makova. 2011. “Genome Analyses Substantiate Male Mutation Bias in Many Species.” *BioEssays: News and Reviews in Molecular, Cellular and Developmental Biology* 33 (12): 938–45.

Zhu, Yuan O., Mark L. Siegal, David W. Hall, and Dmitri A. Petrov. 2014. “Precise Estimates of Mutation Rate and Spectrum in Yeast.” *Proceedings of the National Academy of Sciences* 111 (22): E2310–18.

#### 9. Supplementary Tables

**Table S1**: Dataset information for wolf pedigrees.

| Koch et al. (2019) ID | NCBI Accession |
| --- | --- |
| 480M | SRR9095638 |
| 569F | SRR1518530, SRR1518531, SRR1518532 |
| 629M | SRR9095637 |
| 645F | SRR9095636 |
| 694F | SRR9095635 |

**Table S2**: Dataset information for chimpanzee pedigrees.

| Venn et al. (2014) ID | NCBI Accession | Details |
| --- | --- | --- |
| A | ERR466113 | Sire1 (Pedigree 1) |
| B | ERR466114 | Dam1 (Pedigree 1) |
| D | ERR466116 | Offspring2 (Pedigree 1) |
| E | ERR466117 | Offspring3 (Pedigree 1) |
| F | ERR466118 | Offspring4 (Pedigree 1), Dam2 (Pedigree 2) |
| C | ERR466115 | Sire2 (Pedigree 2) |
| G | ERR466119 | Offspring5 (Pedigree 2) |
| H | ERR466120 | Offspring6 (Pedigree 2) |
| I | ERR466121 | Offspring7 (Pedigree 2) |

**Table S3**: Quantiles and mean of GQ (gray) and depth (white) for variant sites in wolf data.

| ID | 0% | 25% | 50% | 75% | 100% | Mean |
| --- | --- | --- | --- | --- | --- | --- |
| 480M | 0 | 45 | 66 | 99 | 99 | 67.43 |
| 569F | 0 | 99 | 99 | 99 | 99 | 92.45 |
| 629M | 0 | 51 | 75 | 99 | 99 | 71.75 |
| 645F | 0 | 48 | 71 | 99 | 99 | 69.76 |
| 694F | 0 | 51 | 72 | 99 | 99 | 70.73 |
| True_positives (all offspring) | 0 | 42 | 69 | 99 | 99 | 85.47 |
| 480M | 0 | 13 | 18 | 22 | 2958 | 17.84 |
| 569F | 0 | 35 | 44 | 57 | 1912 | 46.06 |
| 629M | 0 | 15 | 20 | 24 | 2847 | 19.82 |
| 645F | 0 | 14 | 19 | 23 | 2553 | 18.66 |
| 694F | 0 | 15 | 19 | 23 | 1754 | 19.39 |
| True_positives (all offspring) | 15 | 19 | 24 | 39 | 49 | 28.11 |

**Table S4**: Quantiles and mean of GQ (gray) and depth (white) for variant sites in chimpanzee data. Chimpanzee F was included in both pedigrees and the values differ slightly between each run. Chimpanzee F values are given as ‘pedigree1/pedigree2’.

| Pedigree | ID | 0% | 25% | 50% | 75% | 100% | Mean |
| --- | --- | --- | --- | --- | --- | --- | --- |
| Pedigree1 | A | 0 | 96 | 99 | 99 | 99 | 90.97 |
|  | B | 0 | 93 | 99 | 99 | 99 | 90.31 |
|  | D | 0 | 72 | 99 | 99 | 99 | 82.79 |
|  | E | 0 | 51 | 75 | 99 | 99 | 72.29 |
| Both | F | 0/0 | 60/60 | 96/96 | 99/99 | 99/99 | 79.41/79.70 |
| Pedigree2 | C | 0 | 54 | 81 | 99 | 99 | 75.10 |
|  | G | 0 | 54 | 81 | 99 | 99 | 73.74 |
|  | H | 0 | 54 | 81 | 99 | 99 | 74.32 |
|  | I | 0 | 60 | 96 | 99 | 99 | 77.94 |
| Pedigree1 | A | 0 | 29 | 36 | 41 | 4369 | 35.61 |
|  | B | 0 | 29 | 36 | 41 | 4700 | 36.70 |
|  | D | 0 | 21 | 26 | 32 | 4699 | 27.76 |
|  | E | 0 | 15 | 19 | 22 | 3373 | 19.78 |
| Both | F | 0/0 | 19/19 | 23/23 | 28/28 | 3509/3875 | 24.94/25.00 |
| Pedigree2 | G | 0 | 17 | 21 | 25 | 3401 | 21.91 |
|  | H | 0 | 16 | 20 | 24 | 3430 | 20.95 |
|  | I | 0 | 16 | 20 | 24 | 4240 | 21.24 |
|  | J | 0 | 18 | 22 | 28 | 4541 | 24.14 |

**Table S5**: Number of sites retained after each RatesTools filtration step in RatesTools for the wolf dataset. ‘Site-filtered sites’ indicates the number of sites after site-level filters have been applied. ‘Region-filtered sites’ is the number of sites after low-quality regions have been removed; this value is the estimate of the ‘callable genome’ length. The mutation rate is the raw estimate from RatesTools, without adjustment for either false positives or false negatives.

| Individual | 629M | 645F | 694F |
| --- | --- | --- | --- |
| All Sites | 2,410,249,677 | 2,410,249,677 | 2,410,249,677 |
| Autosomal Sites | 2,203,158,627 | 2,203,158,627 | 2,203,158,627 |
| Site-Filtered Sites | 1,522,555,977 | 1,470,968,692 | 1,516,943,821 |
| Region-Filtered Sites | 1,278,435,068 | 1,238,792,041 | 1,274,022,488 |
| Candidate DNMs | 94 | 104 | 119 |
| Mutation Rate (95% C.I.) | 3.7e-8 (3.1e-8–4.1e-8) | 4.2e-8 (4.1e-8–3.6e-8) | 4.7e-8 (3.9e-8–5.0e-8) |

**Table S6**: Number of sites retained after each RatesTools filtration step for the chimpanzee pedigree 1 dataset. ‘Site-filtered sites’ indicates the number of sites after site-level filters have been applied. ‘Region-filtered sites’ is the number of sites after low-quality regions have been removed; this value is the estimate of the ‘callable genome’ length. The mutation rate is the raw estimate from RatesTools, without adjustment for either false positives or false negatives.

| Step | D | E | F |
| --- | --- | --- | --- |
| All Sites | 3,230,777,857 | 3,230,777,857 | 3,230,777,857 |
| Autosomal Sites | 2,785,032,019 | 2,785,032,019 | 2,785,032,019 |
| Site-Filtered Sites | 2,368,279,036 | 1,988,140,537 | 2,311,525,550 |
| Region-Filtered Sites | 1,650,850,172 | 1,512,212,130 | 1,627,946,267 |
| Candidate DNMs | 195 | 154 | 192 |
| Mutation Rate (95% C.I.) | 5.5e-8 (5.2e-8–6.1e-8) | 5.1e-8 (4.4e-8–5.4e-8) | 5.6e-8 (5.1e-8–5.9e-8) |

**Table S7**: Number of sites retained after each RatesTools filtration step for the chimpanzee pedigree 2 dataset. ‘Site-filtered sites’ indicates the number of sites after site-level filters have been applied. ‘Region-filtered sites’ is the number of sites after low-quality regions have been removed; this value is the estimate of the ‘callable genome’ length. The mutation rate is the raw estimate from RatesTools, without adjustment for either false positives or false negatives.

| Step | G | H | I |
| --- | --- | --- | --- |
| All Sites | 3,230,766,519 | 3,230,766,519 | 3,230,766,519 |
| Autosomal Sites | 2,785,029,615 | 2,785,029,615 | 2,785,029,615 |
| Site-Filtered Sites | 1,871,482,093 | 1,895,866,311 | 1,982,843,403 |
| Region-Filtered Sites | 1,430,465,769 | 1,446,895,694 | 1,500,689,844 |
| Candidate DNMs | 47 | 79 | 152 |
| Mutation Rate (95% C.I.) | 1.6e-8 (1.4e-8–1.8e-8) | 2.8e-8 (2.4-e8–2.9e-8) | 5.1e-8 (4.5e-8–5.6e-8) |

**Table S8**: Comparison of candidate DNMs identified by RatesTools and those identified by Koch et al. (2019) in the wolf pedigree. ‘Proportion Verified’ gives the fraction of DNMs verified via Sanger sequencing identified in RatesTools. ‘Proportion False Positive’ gives the fraction of DNM candidates shown to be false positive via Sanger sequencing identified as DNM candidates by RatesTools. ‘Proportion failure’ gives the fraction of DNM candidates where Sanger sequencing verification failed. These last three statistics are given as ‘number of RatesTools sites/number of Koch et al. sites’.

| **Individual** | **Koch et al. Total Candidate Sites** | **RatesTools Total Candidate Sites** | **Proportion Verified** | **Proportion False Positive** | **Proportion Failure** |
| --- | --- | --- | --- | --- | --- |
| 629M | 109 | 119 | 5/5 | 1/9 | 0/1 |
| 645F | 106 | 104 | 7/7 | 2/8 | 2/3 |
| 694F | 108 | 94 | 3/3 | 0/12 | 3/4 |

**Table S9**: Comparison of chimpanzee DNMs verified by Venn et al. (2014) to RatesTools results. ‘Overlapping’ gives the number of Venn et al.-verified DNMs found with RatesTools. ‘Multiple Hits’ lists the number of verified Venn et al. DNMs that do not map uniquely to chimpanzee assembly 2.1.4. ‘Non-SNP’ lists the number of confirmed DNMs that were either not genotyped (i.e. no reads mapped to the region) or were not found to be a SNP site. ‘Site-Filtered’ and ‘Region-Filtered’ give the numbers of Venn et al.-verified sites that failed site-level and low-quality-region filters, respectively. ‘DNM Discarded’ lists the number of Venn et al. DNMs that were discarded during RatesTools DNM identification and filtration. For chimpanzee F, one site (denoted by *) was not found to be SNP by RatesTools and was also excluded by the low-quality region filter. Another (denoted by ^) was missing a parental genotype along with exclusion by the region filter.

| **ID** | **Venn et al. verifiDNMs** | **Overlapping** | **Multiple Hits** | **Non-SNP** | **Site-Filtered** | **Region-Filtered** | **DNM Discarded** |
| --- | --- | --- | --- | --- | --- | --- | --- |
| E | 35 | 24 | 5 | 2 | 4 | 1 | 0 |
| F | 25 | 14 | 1 | 2* | 3^ | 6*^ | 1 |

**Table S10**: Number of autosomal candidate single-forward DNMs retained after each filter in the wolf dataset. Results are given as ‘Single-forward mutations (Double-forward mutations:Back mutations)’.

| Individual | 629M | 645F | 694F |
| --- | --- | --- | --- |
| Autosomal Candidates | 695,497 (1,356,316/1,756,466) | 633,719 (1,413,788:1,795,747) | 672,370 (1,336,580:1,759,156) |
| Site Filtration | 32,260 (2,284:14,605) | 24,825 (2,284:13,389) | 25,102 (2,734:14,346) |
| Region Filtration | 1,347 (0:291) | 815 (2:239) | 794 (0:235) |
| Parental Homozygosity Filter | 1,347 (0:150) | 815 (2:142) | 794 (0:135) |
| Koch et al. DNp Filter | 1,057 (0:0) | 564 (2:0) | 541 (0:0) |
| Allele Count Filters | 94 (8:0) | 104 (8:0) | 119 (14:0) |

#### 10. Supplementary Figures

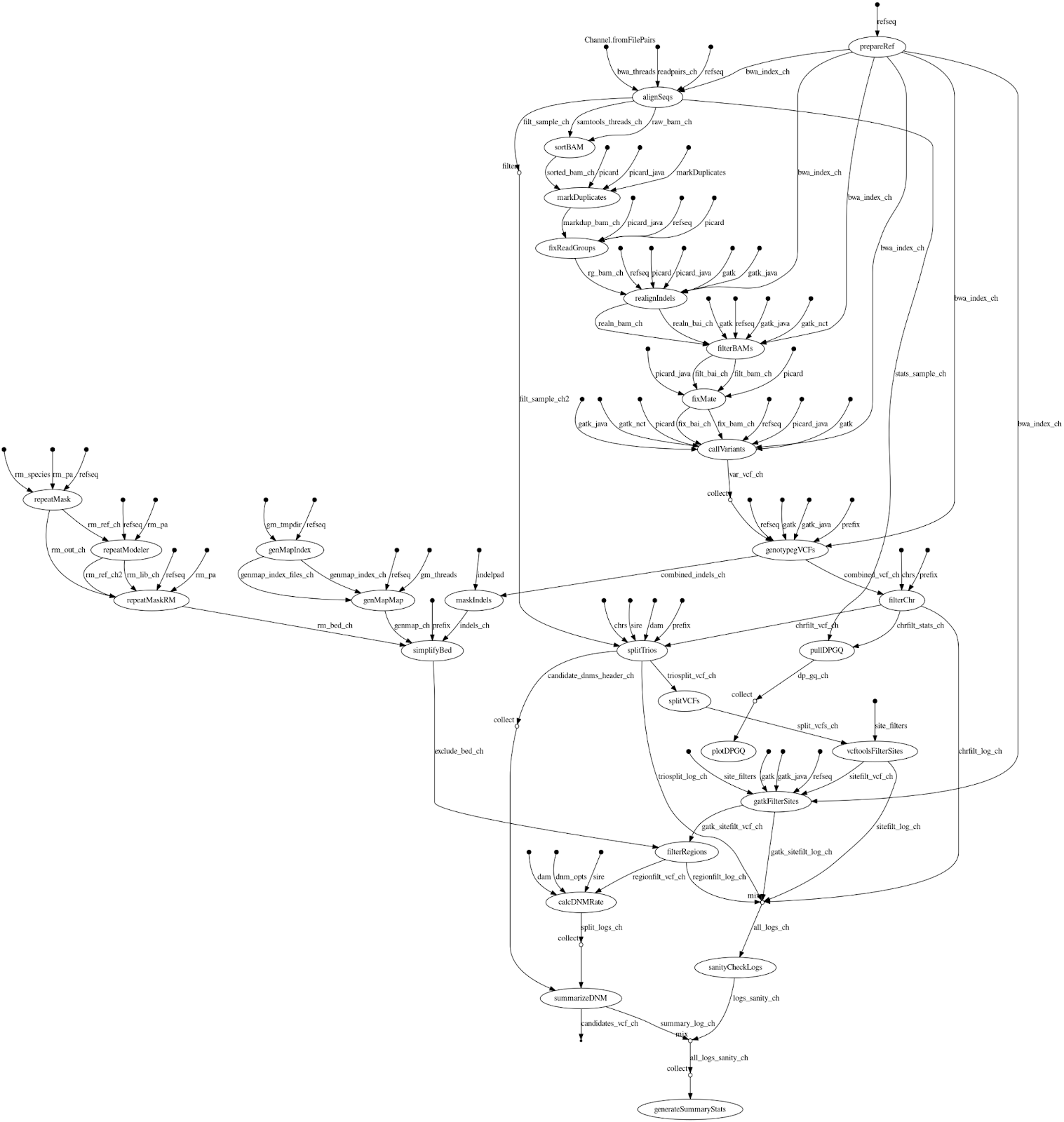

**Figure S1**: Complete directed acyclic graph of the RatesTools pipeline showing all inputs and parameters.

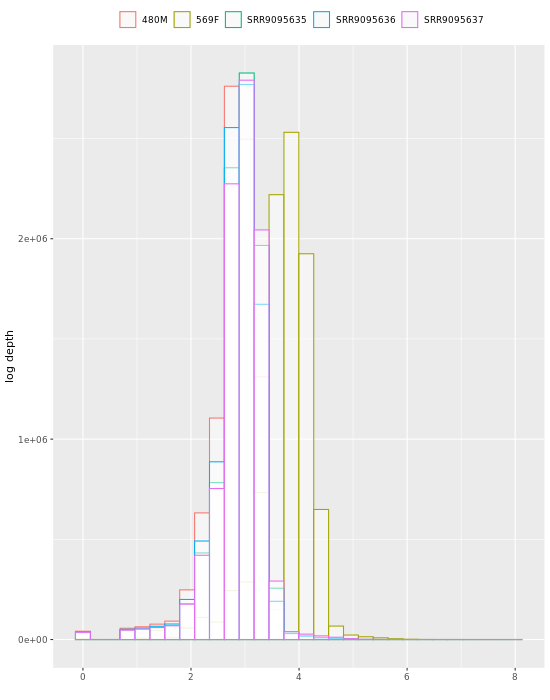

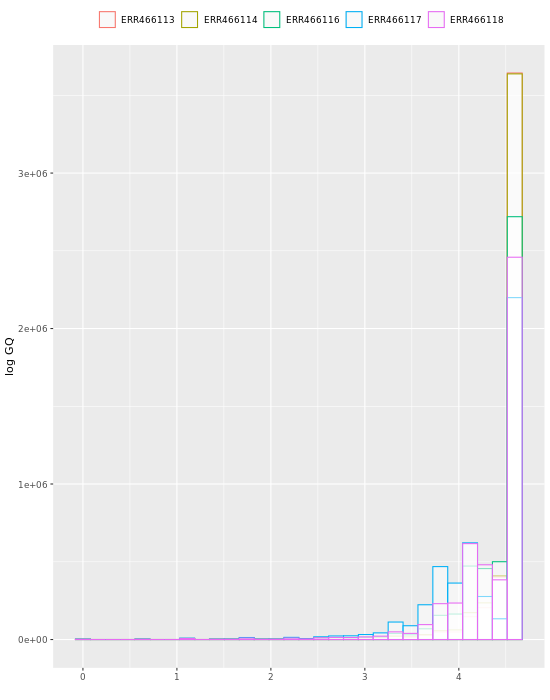

**Figure S2**: Log depth and GQ distributions of variants from pedigreed wolves.

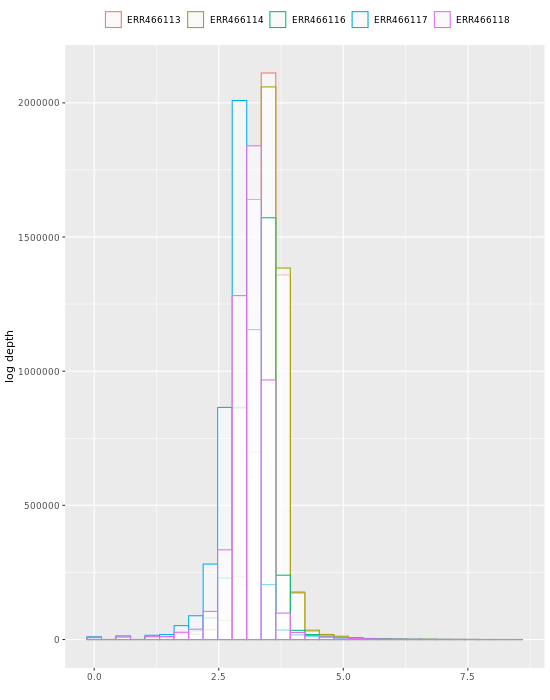

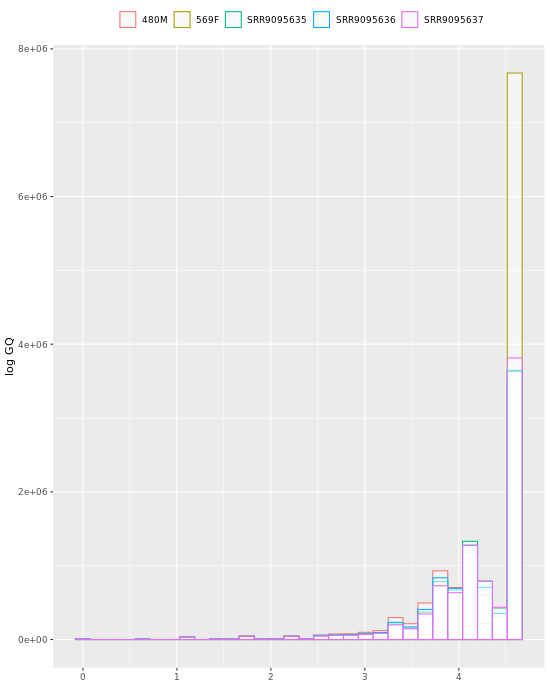

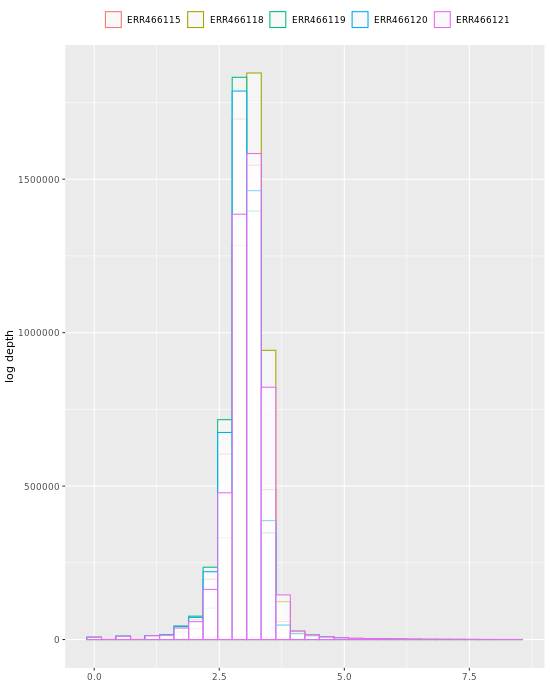

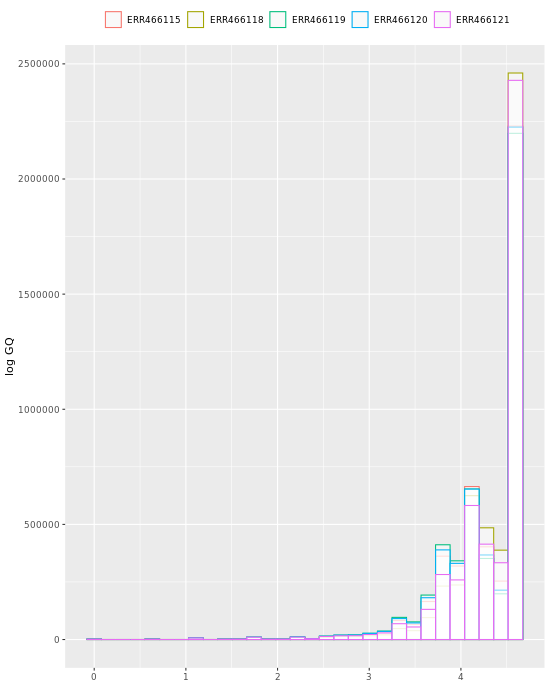

**Figure S3:** GQ and depth distributions for RatesTools candidate *de novo* mutations.
